# A microtissue-based retinal fibrosis platform for drug efficacy testing

**DOI:** 10.64898/2026.06.13.732056

**Authors:** Thomas Hofstetter, Christophe Roubeix, Agnieszka Pawlowska, Peter D. Westenskow, David A. Fluri

## Abstract

**PURPOSE:** Development of a microtissue-based phenotypic screening platform to assess the potency of antifibrotic drugs for patients with the neovascular form of age-related macular degeneration.

**METHODS:** A robust and scalable three-dimensional in vitro model based on primary retinal pigment epithelium (RPE) cells was developed, and a fibrotic disease phenotype was induced. The endpoints included scalable image-based segmentation and quantification of collagen I and fibronectin, bright-field analysis of phenotypic morphological changes, and the analysis of secreted procollagen I levels alongside whole-transcriptome gene expression profiling to demonstrate the potency of compounds in repressing the fibrotic phenotype.

**RESULTS:** The developed model shows similarity to in vivo tissue structures. The three-dimensional constructs form a prominent, polarized monolayer at the periphery. Cellular markers, including Ezrin and ZO-1, confirm epithelial identity and a strongly polarized morphology with junctional structures. Transcriptomic analysis over the culture period demonstrates progressive microtissue maturation.

Pathway modulators induced epithelial-to-mesenchymal transition (EMT) and fibrotic phenotypes. Transcriptomic analysis demonstrated strong marker upregulation. The fibrotic phenotype and its repression by specific small molecule inhibitors were confirmed by measuring secreted procollagen I, quantifying fibronectin and collagen I, and assessing morphological changes via bright-field imaging.

**CONCLUSIONS:** The developed test system exhibits tissue-specific morphology and functionality, showing a high degree of retinal identity. Disease induction led to broad induction of EMT and fibrotic markers, rendering the test system amenable to testing compounds that inhibit or repress fibrotic phenotypes. Production, culture, and endpoint assessment on the Akura platform render the system fully automation-compatible and scalable to higher throughput.

## Introduction

Age-related macular degeneration (AMD) is a leading cause of irreversible vision loss in the elderly. The disease is broadly classified into dry AMD and neovascular (wet) forms. With population aging, the number of affected individuals is projected to increase worldwide to more than 280 million within the next 15 years, posing a significant public health challenge (Wong et al., 2014).

The retinal pigment epithelium (RPE), a key component of the outer blood-retina barrier, plays a central role in the pathophysiology of neovascular AMD. Fibrotic remodeling driven by epithelial-to-mesenchymal transition (EMT) of retinal cells has emerged as a critical step in disease progression (Ishikawa et al., 2016; Little et al., 2018; Tamiya et al., 2010). RPE cells form a highly polarized epithelial monolayer between photoreceptors and the choroid, where they maintain retinal homeostasis by processing metabolic waste, maintaining barrier function, and regulating molecular transport (Maminishkis et al., 2006).

Despite advances in understanding disease mechanisms, modelling fibrosis in neovascular AMD remains challenging. Animal models are often not suitable, as the crucial central part of the retina, the macula, is absent in most commonly used laboratory animals and is found only in humans and non-human primates (S. Chen et al., 2014; Marmorstein & Marmorstein, 2007; Zeiss, 2010). Furthermore, in vivo subretinal fibrosis models are generally induced acutely, exhibit variable phenotypes, and heal spontaneously (Pennesi et al., 2012).

Human *in vitro* systems that recapitulate key pathological features have therefore become essential tools in preclinical research.

Many aspects of retinal dysfunction have been modelled using simple systems based on cell lines, immortalized primary cells, or differentiated pluripotent cells, mainly in two-dimensional formats (Bharti et al., 2022; Dunn et al., 1996). A prominent example is the ARPE-19 cell line which is widely utilized in retinal research due to its robust expansion capacity and ability to form polarized monolayers in transwell assays. However, the translational utility of ARPE-19 cells is constrained by intrinsic genomic instability and the marked underexpression of multiple hallmark retinal genes compared to primary RPE cells (Samuel et al., 2017; Strunnikova et al., 2010).

More advanced systems, including three-dimensional aggregates, retinal organoids or microfluidic systems often suffer from limited reproducibility, scalability, and compatibility with high throughput applications (Bharti et al., 2022; Harkin et al., 2024; Sato et al., 2013; Song et al., 2023; Spirig & Renner, 2024; Usui et al., 2019; Wahle et al., 2023; Zhao & Yan, 2024).

Three-dimensional microtissue constructs generated from primary cells offer a promising intermediate approach, combining physiological relevance with improved experimental robustness and scalability.

Cultured RPE cells have been shown to undergo epithelial-to-mesenchymal transition (EMT) and adopt a fibrotic phenotype in response to pathway modulators including TGF-β, IL-6, PDGF, and TNF-α in multiple in vitro models (Boles et al., 2020; Schiff et al., 2019).

Here, we present a screening-compatible three-dimensional RPE microtissue (hRPE-MT) model based on primary cells. These spheroidal constructs recapitulate key morphological and functional features of native tissue, including cellular polarization and junctional organization.

Upon stimulation with the pro-fibrotic and pro-inflammatory factors TGF-β and TNF-α, the model undergoes EMT-associated fibrotic remodeling, characterized by pronounced morphological changes, extracellular matrix (ECM) reorganization, and upregulation of fibrosis markers.

Capitalizing on this inducibility, we established a screening platform for assessing the antifibrotic potency of test compounds. The inducible model system is highly scalable due to matrix-and hydrogel-free aggregation and culture in fully automation-compatible plates with throughput compatible endpoints.

## Materials and methods

### hRPE-MT aggregation and culture

Primary human RPE cells (HRPEpiC, ScienCell) were expanded for 4 days in epithelial cell medium (EpiCM, ScienCell) supplemented with 2% fetal bovine serum on 0.01% poly-L-lysine (Sigma)-coated culture flasks. Cells were then detached and seeded at 2,000 cells per well in hRPE-MT aggregation medium (InSphero) in Akura 96-well spheroid microplates or Akura 384-well ImagePro plates (InSphero). Following aggregation, hRPE microtissues were maintained at 37 °C, 5% CO₂, and 95% humidity in serum-free epithelial cell medium.

### Compounds and cytokines

SB-431542, RepSox, AM580, CCG-1423 and CP-673451 were obtained from TargetMol (USA) and solubilized in 100% dimethyl sulfoxide (VWR Life Science) to prepare stock solutions.

TNF-α (Sigma) and TGF-β2 (PeproTech) were reconstituted in Milli-Q water and subsequently diluted in phosphate buffered saline (Sigma) containing bovine serum albumin (Merck).

### ATP and LDH assays

Intracellular ATP levels were quantified using the CellTiter-Glo® 2.0 Cell Viability Assay (Promega) according to the manufacturer’s instructions. Lactate dehydrogenase (LDH) leakage into the supernatant from damaged cells was measured using the LDH-Glo™ Cytotoxicity assay (Promega) according to the manufacturer’s protocol. Briefly, cell culture supernatants were diluted 1:5 in LDH storage buffer and the diluted samples were mixed 1:1 with detection reagent in white half-area assay plates (Greiner Bio-One). Following incubation for 1h at room temperature on an orbital shaker, luminescence was recorded with a Tecan Spark 10M plate reader.

### Histology

Microtissues from each treatment group were pooled, washed with PBS, and fixed in 4% paraformaldehyde (Alfa Aesar) overnight at 4°C. Following fixation, samples were washed with PBS, pelleted in 1.7% agarose, and processed for paraffin embedding, sectioning, and mounting on poly-L-lysine-coated glass slides by an external service provider (Sophistolab AG).

Immunohistochemical (IHC) and histochemical (HC) stainings were performed at Sophistolab AG using the following primary antibodies: ezrin (Abcam, Cat. No.: ab4069), zo-1 (Thermo Fisher, Cat. No.: 33-9100), collagen I (Abcam, Cat. No.: ab88147), fibronectin (Santa Cruz, Cat. No.: sc-59826) and α-smooth muscle actin (Sigma, Cat. No.: 202M-94). Images were acquired at 40x magnification using a Leica DMi8 microscope equipped with a DMC4500 digital camera.

### RNA sequencing and analysis

RNA sequencing was performed by BioSpyder Technologies, Inc. using TempO-Seq™ templated oligo-sequencing technology with the Human Whole Transcriptome assay. Following sample collection, hRPE-MTs were washed with calcium-and magnesium-free PBS and lysed in 15 μL of 1× Enhanced Lysis Buffer (BioSpyder Technologies, Inc.). Lysates were shipped to BioSpyder Technologies for library preparation and sequencing on an Illumina platform. Following sample demultiplexing and generation of FASTQ files, read alignment and count generation were performed using the TempO-SeqR data analysis platform provided by BioSpyder Technologies. Downstream analysis of TempO-Seq data was performed by InSphero using a proprietary analytical pipeline implemented in the R programming language. Probe-level raw counts were collapsed to gene-level counts by summing counts from probes associated with the same gene. Data normalization, principal component analysis (PCA), and differential expression analysis (DEA) were performed using the DESeq2 R package. Surrogate variable analysis (SVA) was applied using the sva R package to account for unwanted sources of variation and unintended batch effects. Pre-ranked gene set enrichment analysis (GSEA) was performed using the clusterProfiler R package. Genes were ranked according to log₂ fold change (log2FC) values derived from the DEA output, and the normalized enrichment score (NES) was used as the enrichment metric. The gene set universe used for GSEA comprised selected subcollections from the Molecular Signatures Database (MSigDB), version 2025.1, including Hallmark, PID, BioCarta, Reactome, and WikiPathways gene sets. For both DEA and GSEA, p values were adjusted for multiple testing using the Benjamini-Hochberg false discovery rate (FDR) procedure. Applicable FDR thresholds are reported together with the corresponding results.

### Concentration range finding

For cytotoxicity assessment, hRPE microtissues were cultured for 10 days with medium changes on days 0, 3, 5, and 7. Compounds were applied in half-log dilution series (50-0.05 μM) on days 0, 3, 5, 7 following medium exchange using a TECAN D300e Digital Dispenser. All conditions were normalized to 0.1% DMSO. Supernatants were collected prior to each medium change for LDH quantification. Intracellular ATP levels were measured on day 10.

### Ocular Fibrosis Induction

For efficacy assessment, human retinal pigment epithelium microtissues (hRPE-MTs) were cultured for 10 days in serum-free epithelial cell medium with medium changes on days 0, 3, 5, and 7. Fibrosis was induced by treatment with 15 ng/mL TNF-α and 15 ng/mL TGF-β2 for 72h starting on day 0, followed by a secondary pulse of 5ng/mL TNF-α and 5 ng/mL TGF-β2 on day 3 for an additional 48 h.

Compounds were applied on days 0, 3, 5, and 7 following medium exchange using a TECAN D300e Digital Dispenser and tested at two concentrations based on prior range-finding experiments. All conditions were normalized to 0.1% DMSO. Supernatants were collected prior to each medium change for secreted pro-collagen I quantification. On day 10, intracellular ATP levels were measured, and microtissues were processed for high-content imaging (HCI).

### Pro-collagen-I assay

Secreted pro-collagen-I levels in supernatants collected on days 3, 5, 7, and 10 were quantified using an HTRF® Human Pro-Collagen Type 1 Detection kit (Revvity) according to the manufacturer’s instructions. Fluorescence was measured using a Tecan Spark 10M plate reader.

### High content imaging

On day 10, hRPE-MTs were washed once with PBS and fixed with 4% paraformaldehyde for 15 min in Akura 384 well plates. Following three PBS washes, samples were permeabilized with 0.5% Triton X-100 in PBS for 1h at room temperature and blocked in 10% horse serum with 0.2% Triton X-100 in PBS for 1h at RT.

Samples were stained overnight at 4 °C, protected from light, in blocking buffer containing Alexa Fluor 647-conjugated anti-collagen I (Abcam, 1:200), Alexa Fluor 555-conjugated anti-fibronectin (Abcam, 1:50), phalloidin-iFluor 488 (Abcam, 1:4000), and DAPI (Sigma, 1:250). After staining, samples were washed four times with wash buffer (0.2% Triton X-100 in PBS) and maintained in PBS (50 µL/well). Except for fixation, all incubation steps were performed on an orbital shaker (300 rpm).

Imaging was performed using a Yokogawa CQ1 spinning-disc confocal high-content imager using a 20x objective. Z-stacks were acquired from the bottom with a total imaging depth of 80 μm with 5 μm step size.

Image analysis was conducted using CellPathfinder software (Yokogawa). Collagen I and fibronectin were quantified by threshold-based 3D analysis within spheroid masks generated from phalloidin staining. Total signal intensity within this spheroid mask was quantified for both markers. To account for variability in imaging depth caused by light scattering, the data was normalized for each spheroid individually based on nuclei counts determined from the DAPI signal.

Phalloidin was quantified as a texture feature in a single z-plane using the integrated texture metric within binarized regions of interest. All readouts were normalized to fibrotic controls set to 100%.

### Resin embedding and section imaging with SEM

Microtissues were fixed in 2.5% glutaraldehyde (EM grade; Polysciences Europe GmbH, Germany) and 2% formaldehyde (EM grade; Polysciences) in 0.15 M cacodylate buffer supplemented with 2 mM calcium chloride (Merck & Cie, Switzerland). In order to enable safe handling of the samples during the following washing and staining steps, the microtissues were embedded in freshly prepared low gelling temperature agarose (4%; Carl Roth GmbH, Germany). After gelling on ice, small cubes containing one microtissue each were cut from the agarose block and washed three times in cacodylate buffer. Samples were stained according to published protocols (Deerinck et al., 2022). Briefly, samples were postfixed with 2% osmium tetroxide (Polysciences) supplemented with 1.5% potassium ferrocyanide (Merck & Cie) and 2mM calcium chloride, followed by 1% thiocarbohydrazide (Sigma-Aldrich, Deutschland), 2% osmium tetroxide, 1% aqueous uranyl acetate (Polysciences) and Walton’s lead aspartate.

Between the staining steps, samples were washed in double distilled water. Then, the samples were dehydrated in increasing concentrations of ethanol, washed in acetone and infiltrated with Epon using a graded series of resin concentrations (2 × 25%, 2 × 50%, 2 × 75%; Epoxy Embedding Kit, Sigma-Aldrich, Germany). Samples were kept in 75% Epon in acetone overnight in the fridge. The following day, samples were infiltrated 3x with fresh Epon for 1h, 2h, and 3h respectively, at RT on a shaker, then transferred into fresh Epon resin and polymerized at 60°C for 3 days. Except for the lead aspartate step, which was carried out in an oven at 60 °C, all steps until 75% Epon were performed in a BioWave Pro+ Tissue Processor (Ted Pella Inc., USA).

Ultrathin sections (100 nm thickness) were cut with a diamond knife (Diatome Ltd., Switzerland) using a Leica UC7 ultramicrotome (Leica Microsystems, Switzerland). Sections were collected on silicon wafer substrates and imaged with a Zeiss Merlin scanning electron microscope equipped with the ATLAS automated imaging platform (Carl Zeiss Microscopy, Germany). Images were acquired under high-vacuum conditions using the sensBSD backscattered electron detector at an accelerating voltage of 2 kV, a beam current of 200 pA, and a working distance of 5 mm. The final image resolution corresponded to a pixel size of 10 nm.

### Statistical analysis

Data was analyzed and plotted using GraphPad Prism (Dotmatics). Results are presented as mean ± standard deviation (SD), with individual data points representing single hRPE microtissues. Statistical significance was assessed by one-way ANOVA followed by Dunnett’s multiple comparisons test, comparing each group to the vehicle control (range-finding studies) or fibrotic control (efficacy studies).

## Results

### 3D retinal pigment epithelial model

Primary human retinal pigment epithelial (hRPE) cells were aggregated into microtissues in 96-and 384-well ultra-low-attachment plates. Highly consistent (diameter: 220 µm ± 10.7 µm, coefficient of variation < 5%) spheroidal aggregates formed within three days (Figure 1A), after which cultures were transitioned to serum-free maintenance conditions, and maintained for up to 28 days.

**Figure 1.**
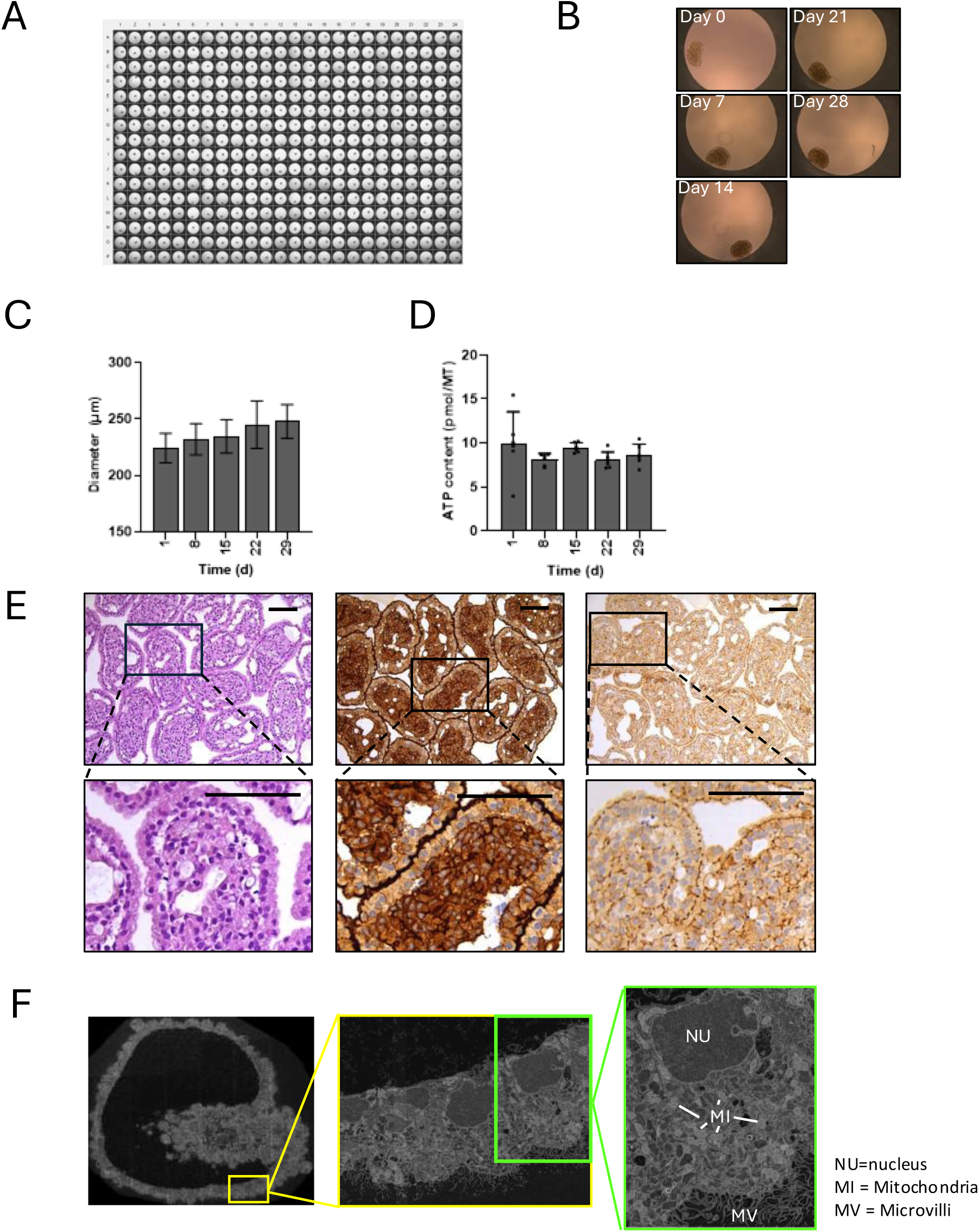
Morphological characterization and microtissue architecture of human retinal pigment epithelium microtissues (hRPE-MTs). (A) Representative brightfield overview scan of hRPE-MTs aggregated in a 384-well plate format. (B) Representative brightfield micrographs illustrating microtissue morphology over the course of culture. Quantification of microtissue diameter (C) and ATP content (D) as a function of culture time. (E) Histological analysis of formalin-fixed, paraffin-embedded (FFPE) and sectioned microtissues stained with hematoxylin and eosin (left), anti-Ezrin antibody (apical marker, middle), and anti-ZO-1 antibody (tight junction marker, right). (F) Ultrastructural characterization by scanning electron microscopy (SEM). Scale bars = 100 µm.

Bright-field micrographs at days 0, 7, 14, 21 and 28 revealed progressive morphological maturation (Figure 1B). A distinct peripheral epithelial monolayer became evident by day 7 and became further pronounced with increasing culture time. By day 14, cystic structures emerged, accompanied by a gradual increase in tissue size through day 28 (Figure 1C).

Microtissue viability was assessed over the four-week culture period by quantifying intracellular ATP levels. No significant differences were observed across time points, indicating stable viability of the 3D RPE constructs (Figure 1D).

For further characterization, microtissues were collected at day 28, paraffin embedded, sectioned, and stained with marker specific antibodies. Hematoxylin and eosin (H&E) staining revealed a well-defined peripheral monolayer of polarized cells, with nuclei localized basolaterally and oriented toward the interior of the spheroid (Figure 1E, left).

Immunostaining for Ezrin, an apically localized microvilli-associated protein, further confirmed the pronounced polarization of the epithelial monolayer (Figure 1E, middle).

In vivo, RPE cells form a highly specialized barrier characterized by well-developed intercellular junctions. Microtissues were analyzed for expression of the tight junction protein zonula occludens-1 (ZO-1). Immunostaining revealed localized ZO-1 expression at cell-cell contacts within the peripheral monolayer, indicating the formation of functional barrier structures in the 3D microtissues (Figure 1E, right).

Ultrastructural analysis by scanning electron microscopy (SEM) supported these findings. Microtissues exhibited a polarized epithelial monolayer surrounding a less organized cellular core with interspersed cyst-like structures (Figure 1F). Apical microvilli were oriented toward the outward-facing surface, and nuclei were oriented basolaterally in the epithelial monolayer. Cells displayed abundant mitochondria and vesicular structures, consistent with high metabolic activity.

To characterize the model at the transcriptional level, whole-transcriptome profiling was performed on RPE microtissues at days 0, 15, and 29 using TempO-Seq. Principal component analysis revealed clear segregation of samples by culture time along principal component 1 (PC1) (Figure 2A), with tight clustering of biological replicates.

**Figure 2.**
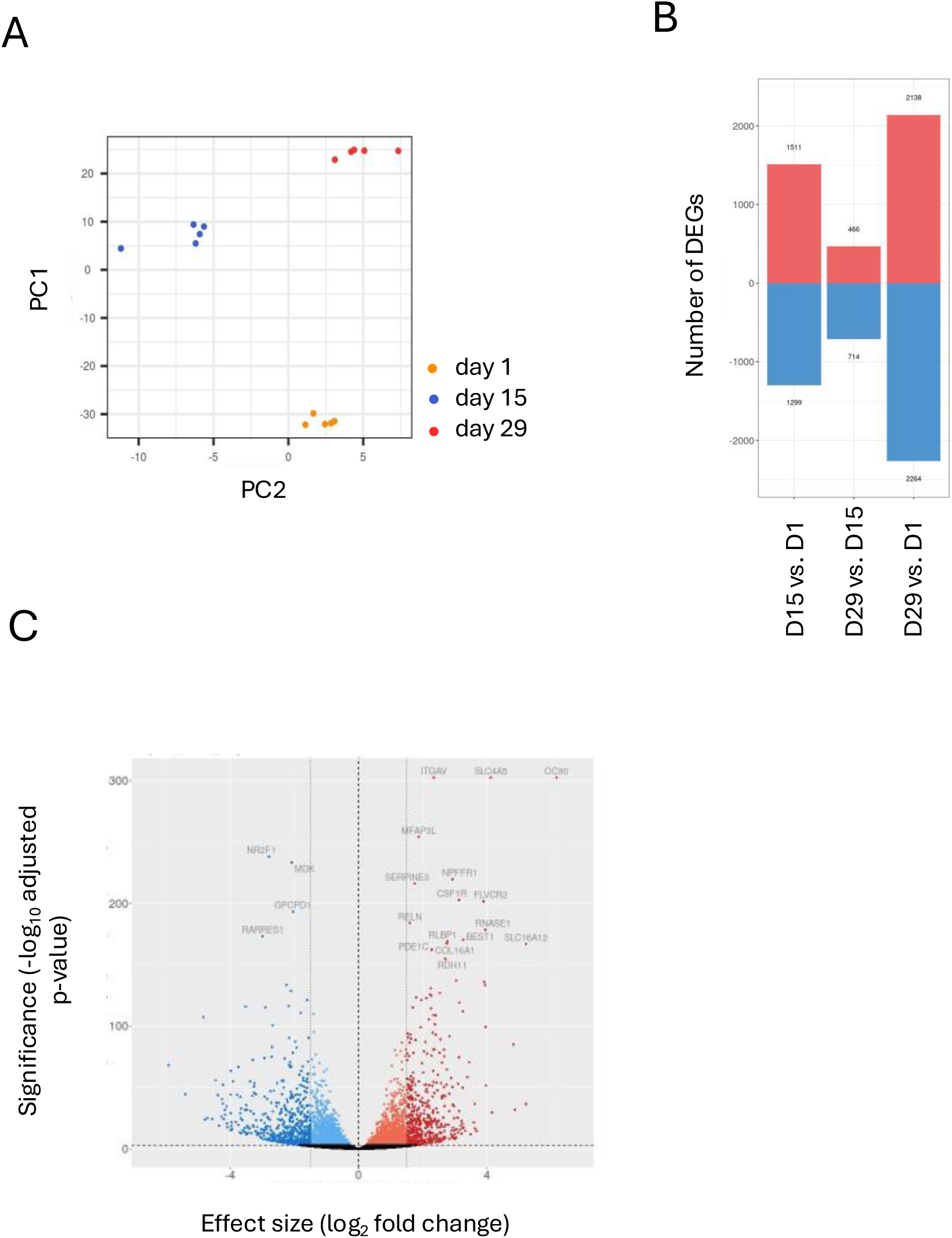
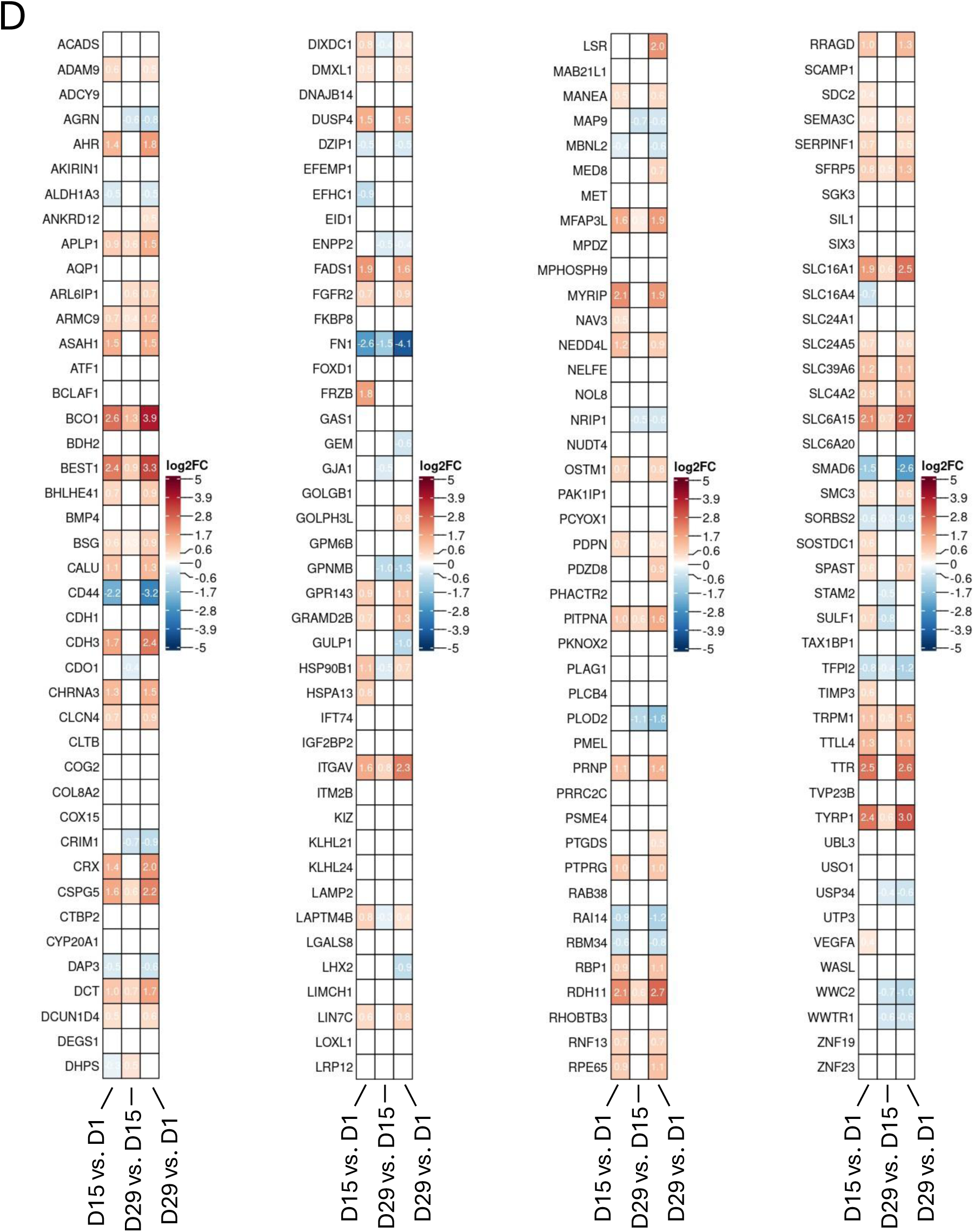
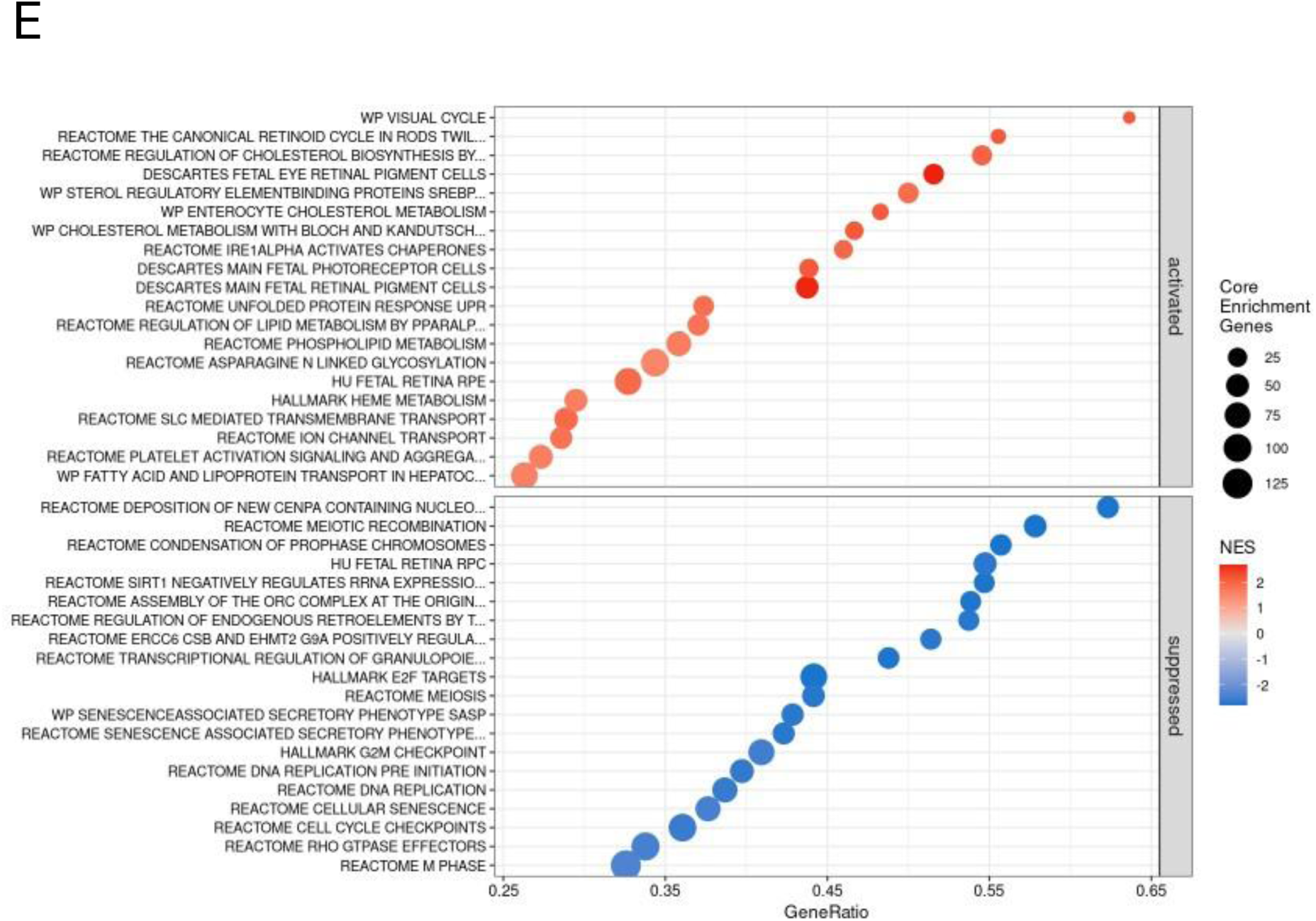
**Whole-transcriptome analysis of human retinal pigment epithelium microtissues (hRPE-MTs) over the course of culture. (**A) Principal component analysis (PCA) of microtissues collected at three culture timepoints, illustrating global transcriptional differences over time. (B) Summary of differentially expressed genes (DEGs) identified between culture timepoints (FDR < 0.001). (C) Volcano plot showing differential gene expression between days 29 and day 1 (FDR < 0.001). (D) Heatmap depicting log2FC values for a panel of 168 retinal identity genes across time points. Columns represent pairwise comparisons: day 15 vs. day 1 (left), day 29 vs. day 15 (middle), and day 29 vs. day 1 (right). Red indicates upregulation, blue indicates downregulation and white indicates non-significant change (FDR < 0.001). (E) Gene set enrichment analysis highlighting the top 20 significantly enriched gene sets among upregulated and downregulated genes between day 29 and day 1 (FDR < 0.05).

DEA revealed transcriptional remodeling, with more than 2,000 genes significantly up-or downregulated between days 0 and 29 (FDR < 0.001). Comparisons between days 0-15 and days 15-29 showed a reduction in differential expression over time, indicating progressive stabilization of the transcriptional profile toward a more mature state (Figure 2B).

Hallmark RPE genes, including retinaldehyde-binding protein 1 (RLBP 1), bestrophin 1 (BEST1), collagen type XVIII alpha 1 chain (COL18A1), solute carrier family 16 member 12 (SLC16A12), and retinol dehydrogenase 11 (RDH11), were among the most strongly induced transcripts during microtissue maturation in 3D (Figure 2C).

To further assess maturation in hRPR microtissues, we analyzed a panel of 168 genes, primarily based on a previously defined retinal gene set (Strunnikova et al., 2010), across 29 days of culture. A substantial proportion of these (42%) showed transcriptional upregulation over time.

Consistent with whole-transcriptome analysis, this retinal gene subset exhibited more pronounced activation between days 0 and 15 compared with days 15 to 29.

Upregulated genes were enriched for key retinal functional categories including visual processes (CRX, RPE65, TRPM1, GPR143 and RDH11), stimulus sensing and signal transduction (AHR, CHRNA3, FGFR2, FRZB, SFRP5, SostDC1, VEGFA, SEMA3C, PTPRG), pigment metabolism (DCT, TYRP1, GPR143, MYRIP), retinoid metabolism and phototransduction (CRX, RPE65, SFRP5, TIMP3, BEST1, RDH11 and RBP1) and transporter activity (SLC4A2, SLC16A1, SLC24A5, SLC6A15, SLC39A6) (Figure 2D).

GSEA revealed robust upregulation of retinal pigment epithelium-specific gene sets during 3D culture maturation. The top 20 enriched gene sets included key functional categories such as the visual and retinoid cycles, retinal pigment epithelial cells, and photoreceptor-associated pathways (Figure 2E). In contrast, the top 20 downregulated gene sets were primarily associated with chromatin remodeling and cell cycle processes, including meiotic recombination, DNA replication, and cell cycle checkpoint regulation.

### Disease induction

Using the human RPE microtissue (hRPE-MT) model, we established an inducible model of ocular fibrosis (Figure 3A). Microtissues were treated with a pro-fibrotic cocktail of TGF-β2 and TNF-α for five days, followed by a five-day period in the absence of inducers prior to analysis.

**Figure 3.**
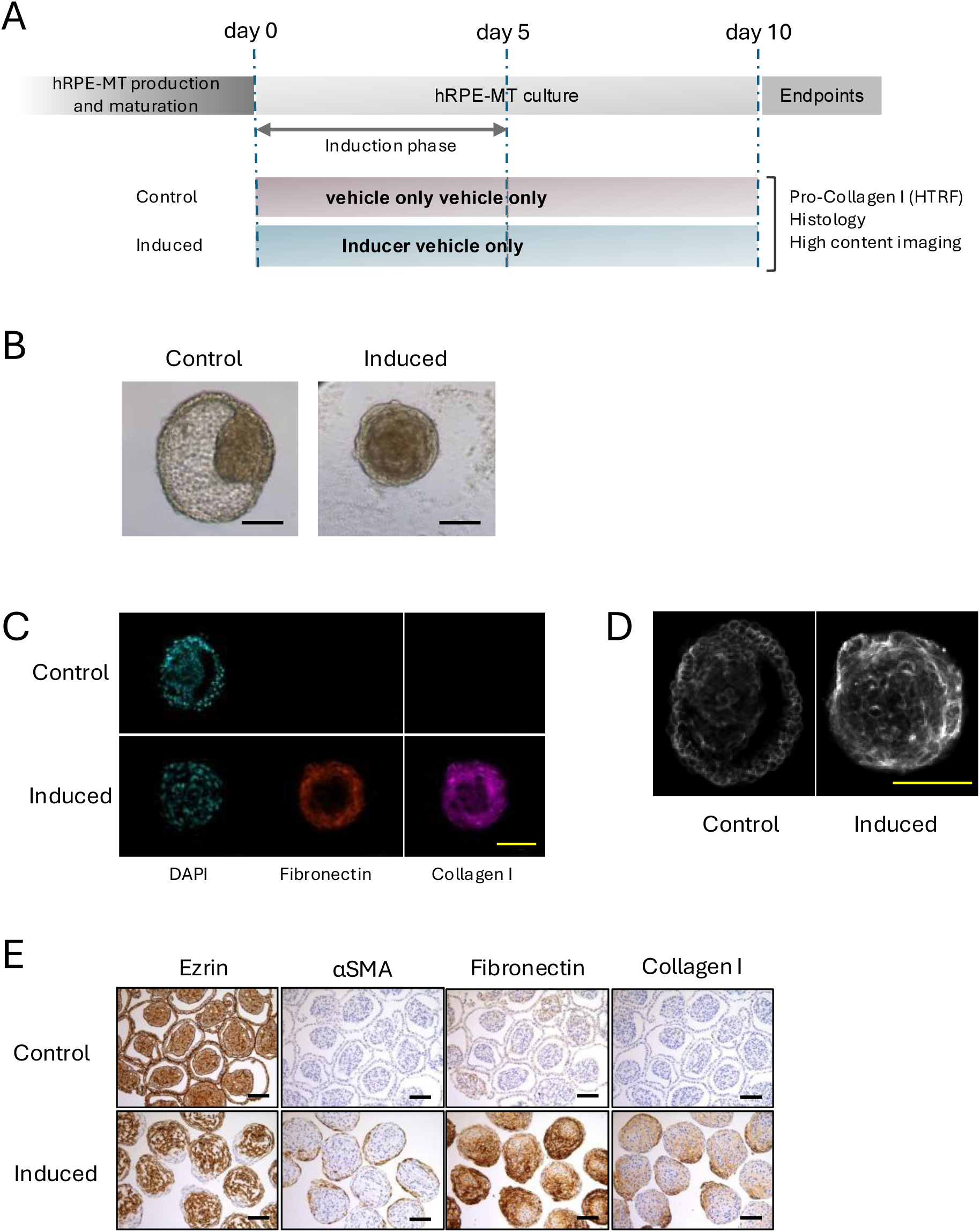
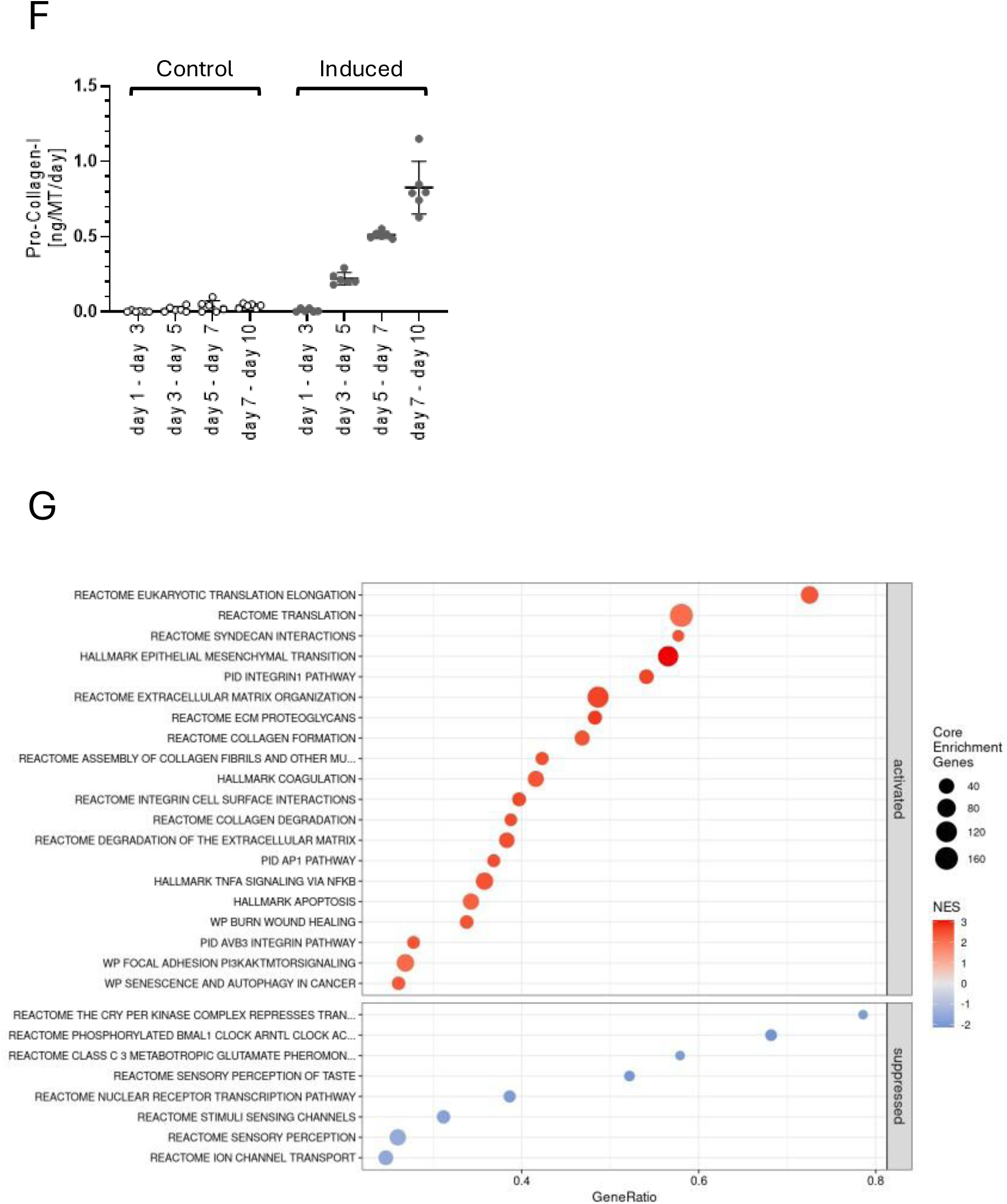
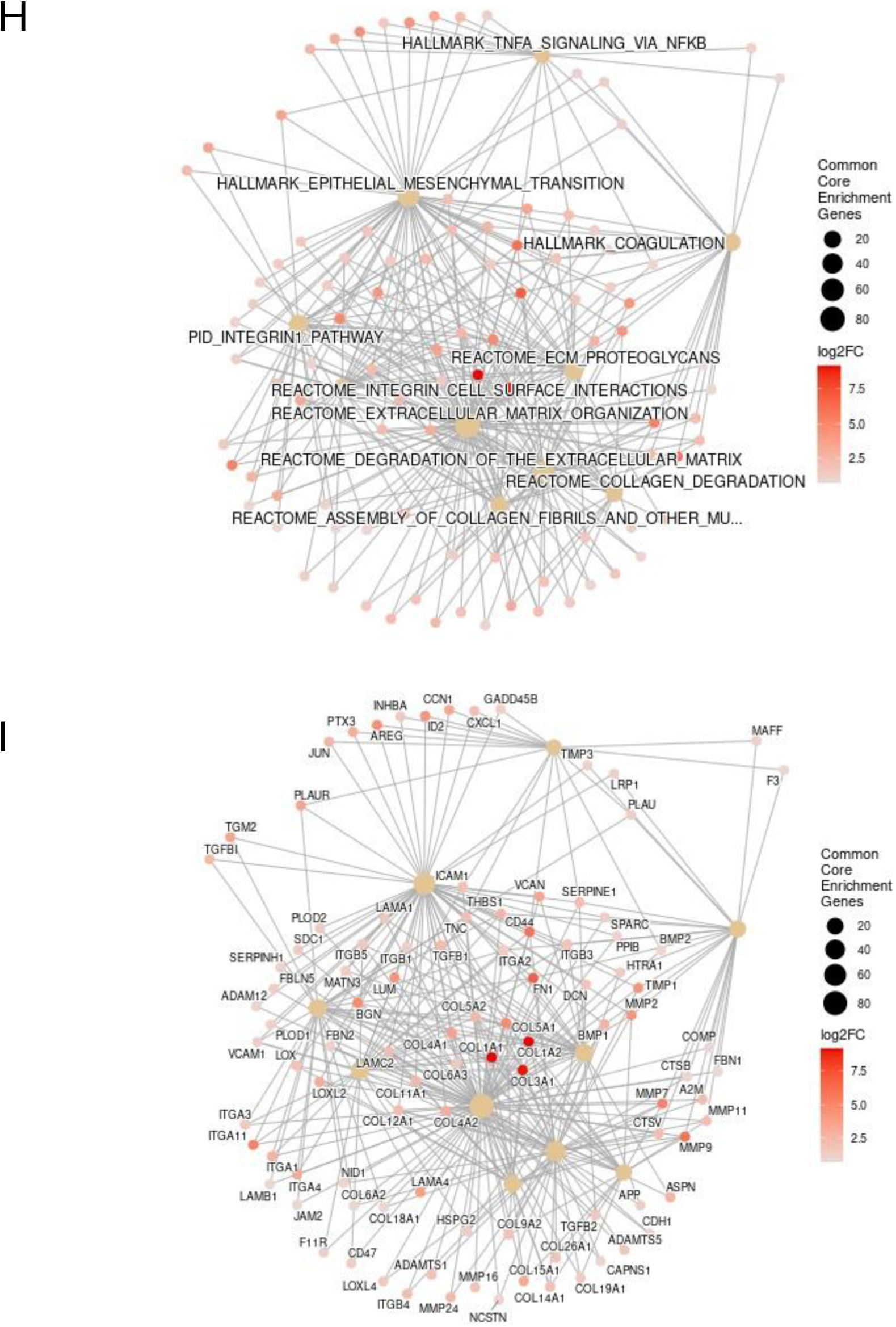
Induction and characterization of fibrotic changes in hRPT-MTs. (A) Schematic overview of the experimental design for fibrosis induction and downstream assays. (B) Representative bright-field micrographs of hRPT-MTs at day 10 post induction, showing uninduced control (left) and induced microtissue (right). (C) Confocal micrographs of induced hRPT-MTs immunostained for fibronectin and collagen I; nuclei are counterstained with DAPI. (D) Confocal visualization of F-actin organization using phalloidin staining in untreated control tissue (left) and induced fibrotic tissue (right). (E) Histological analysis of formalin-fixed, paraffin-embedded (FFPE) and sectioned microtissues stained for Ezrin, α-SMA, fibronectin and collagen I. Untreated controls (top) are compared with induced conditions (bottom). (F) Quantification of secreted pro-collagen I levels in culture supernatants using a homogenous time resolved fluorescence (HTRF) assay, comparing uninduced and induced hRPT-MTs over time. (G) Gene set enrichment analysis (GSEA) identifying the top 20 significantly enriched gene sets among upregulated gene sets and the top 5 among downregulated gene sets in induced versus control conditions (FDR < 0.01). (H, I) Network visualization of common core enrichment genes in top 10 significantly enriched gene sets (FDR < 0.01). Red nodes represent individual genes, with color intensity corresponding to log2FC, while edges indicate functional connectivity. Larger beige nodes denote gene-gene set associations.

Bright-field imaging revealed marked morphological differences between induced and control conditions. Control microtissues displayed heterogenous architectures with dense cellular regions interspersed with cystic structures and a distinct peripheral epithelial monolayer (Figure 3B, left). In contrast, induced microtissues exhibited compaction into dense, highly spherical structures lacking cystic features (Figure 3B, right).

Confocal microscopy-based immunofluorescence analysis demonstrated robust expression of fibrotic markers fibronectin and collagen I in induced microtissues, whereas these markers were undetectable in uninduced controls (Figure 3C).

Phalloidin staining of F-Actin revealed well-organized epithelial architecture in control microtissues (Figure 3D, left). In contrast, induced microtissues exhibited pronounced cytoskeletal remodeling, characterized by loss of epithelial organization, disassembly of actin structures, and the emergence of high-intensity actin fibers consistent with stress fiber formation (Figure 3D, right).

Induced and control microtissues were harvested for histological analysis. Ezrin staining revealed a transition from uniform expression throughout control spheroids to region-specific loss of retinal identity in induced microtissues 10 days post-induction.

Consistent with fibrotic remodeling, collagen-I, fibronectin, and α-smooth muscle actin (α-SMA) were markedly upregulated in induced conditions (Figure 3E).

Distinct spatial expression patterns were observed, with α-SMA localized predominantly at the periphery, collagen I enriched at the poles, and fibronectin distributed throughout the microtissue (Figure 3E, bottom).

The dynamics of EMT and fibrotic remodeling were assessed by quantifying secreted procollagen I (PC-I) levels in culture supernatants over the course of induction and subsequent culture (Figure 3F). PC-I was undetectable in uninduced controls, whereas induced microtissues exhibited a progressive increase in PC-I levels, reaching maximal levels by day 10. Notably, procollagen secretion continued to rise after withdrawal of the inducing stimuli at day 5, indicating sustained fibrotic activity beyond the initial induction phase.

Whole transcriptome profiling by TempO-Seq, followed by gene set enrichment analysis comparing induced and control microtissues at day 10 post-induction, revealed a pronounced fibrotic signature. Among the top 20 enriched pathways, the majority were directly associated with epithelial-to-mesenchymal transition (EMT) and fibrosis (Figure 3G, top), whereas downregulated gene sets were predominantly linked to neuronal function and sensory perception (Figure 3G, bottom).

Network analysis identified a highly interconnected module centered on extracellular matrix organization, linked to EMT, collagen metabolism, and coagulation pathways (Figure 3H). At the gene level, strong induction was observed for multiple members of the collagen, matrix metalloproteinase (MMP), and integrin families, as well as key fibrosis-associated genes including fibronectin, Adam12, CD44 and BGN (Figure 3I).

EMT is a central driver of subretinal fibrosis and is regulated by multiple signaling pathways, most prominently TGF-β, Rho/MRTF/SRF and PDGF signaling. In contrast, pathways such as BMP7 and retinoic acid receptor (RAR/RXR) signaling have been reported to counteract EMT without directly inhibiting these core signaling axes. To evaluate pathway-specific modulation of EMT in hRPE microtissues, we selected a panel of antifibrotic compounds targeting these mechanisms: SB-431542 and RepSox (TGFβ-pathway inhibitors), CP-673451 (PDGFR inhibitor), CCG-1423 (Rho pathway inhibitor) and AM580 (RAR agonist).

The assay design involved co-treatment with pro-fibrotic stimuli and inhibitors during the initial phase, followed by inhibitor-only treatment in a subsequent phase (Figure 4A).

**Figure 4.**
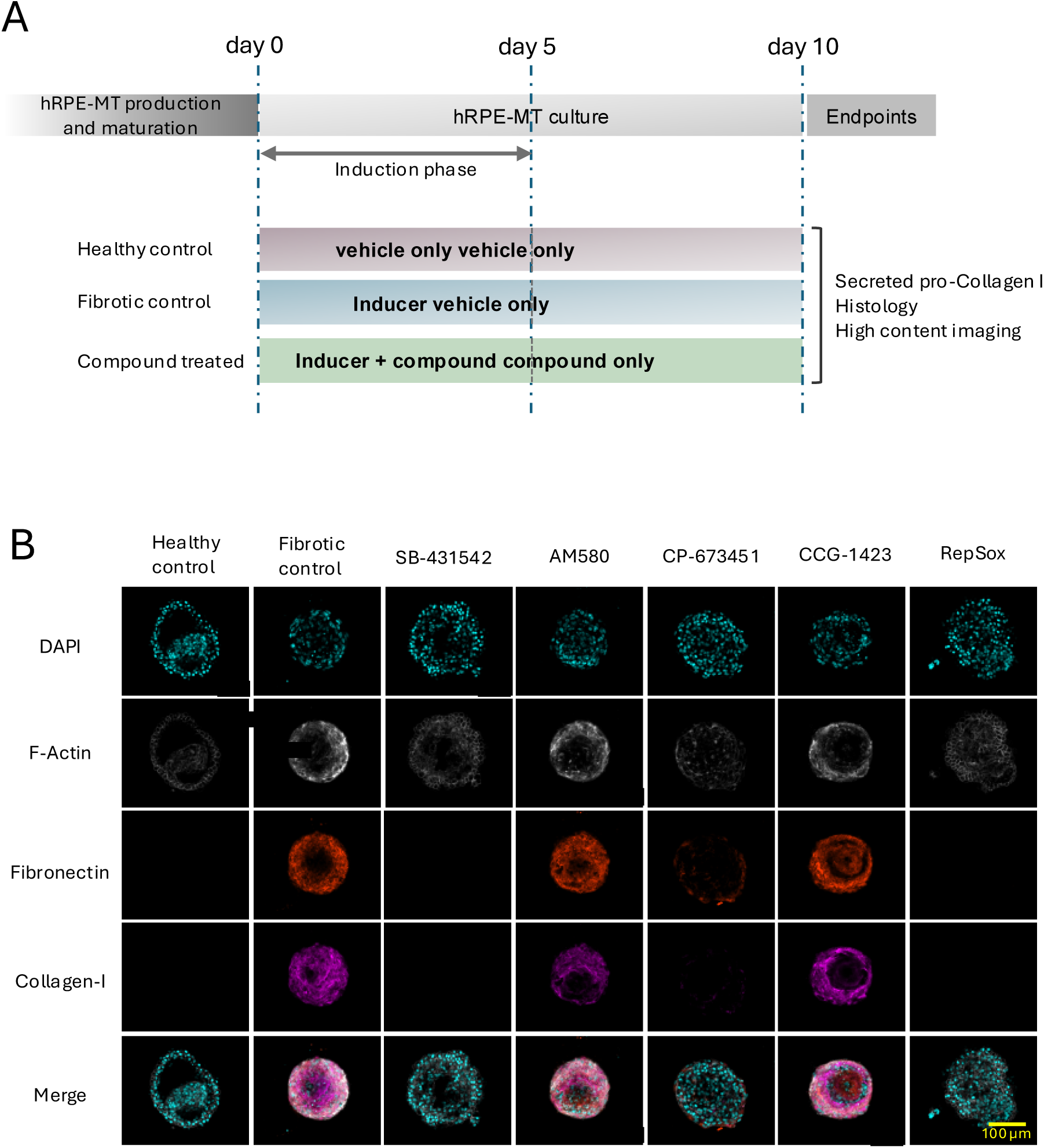
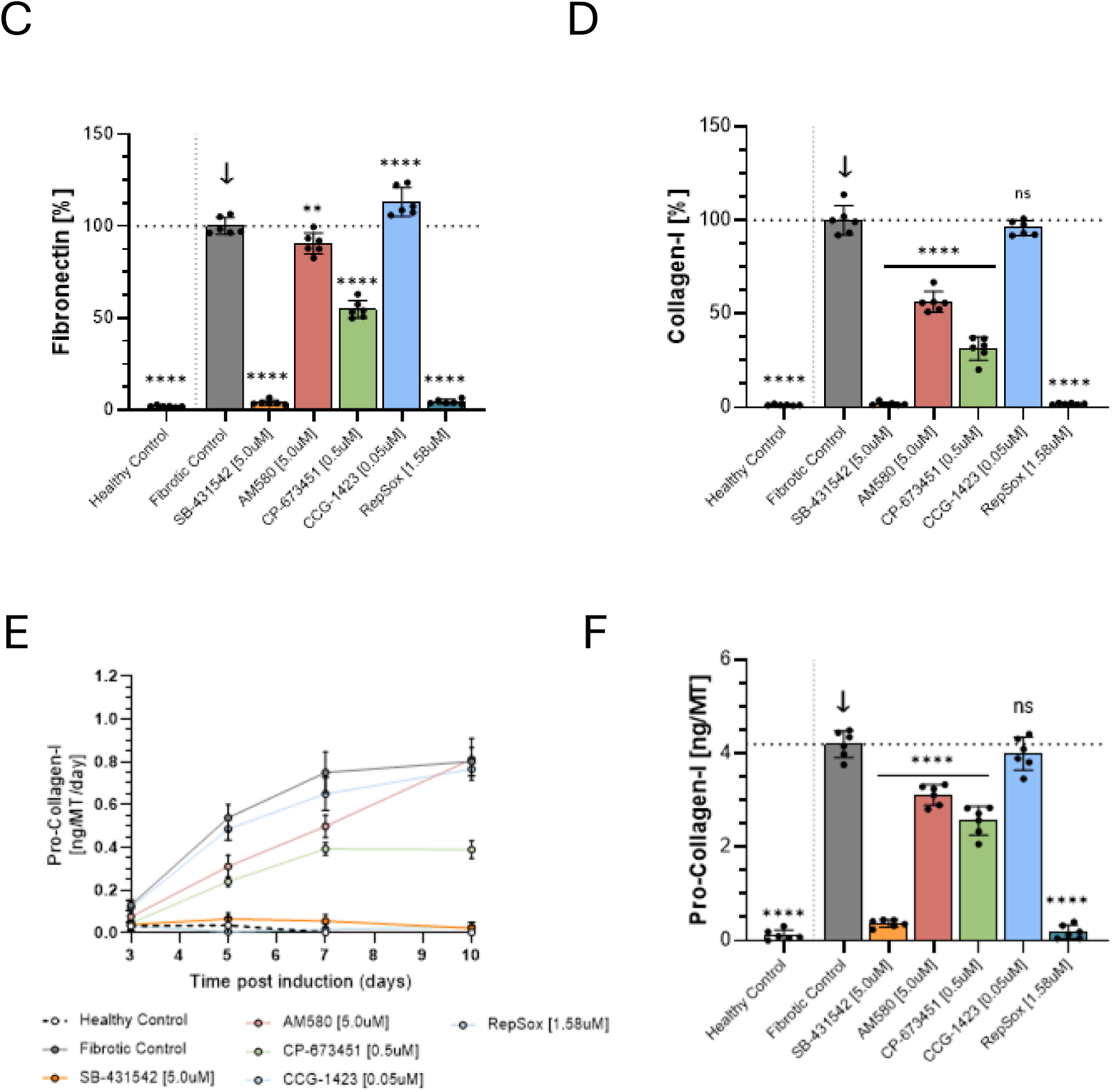
Inhibition of fibrotic phenotypes by pathway-specific inhibitors. (A) Schematic overview of experimental design. (B) Representative confocal immunofluorescence micrographs of control and induced hRPT-MTs cultured in the absence or presence of pathway inhibitors. Microtissues were immunostained for fibronectin and collagen I, with F-actin labeled by phalloidin and nuclei counterstained with DAPI. (C, D) Quantification of fibronectin (C) and collagen I (D) fluorescence intensities in segmented microtissues. Signal intensities in induced tissues without inhibitors were normalized to 100%. (E) Time course of secreted pro-collagen I levels in culture supernatants of hRPT-MTs over the duration of the experiment. (F) Quantitative analysis of cumulative pro-collagen I secretion across the experimental period.

Compounds were first titrated to determine the optimal concentrations for efficacy testing (Figure S1). Subtoxic concentrations were selected based on IC_50_ values and used to define the working concentrations applied in the fibrosis assay.

High content imaging analysis at day 10 post-induction, assessing F-actin, fibronectin, and collagen I, revealed marked differences in antifibrotic efficacy among the tested compounds. Control microtissues exhibited no fibrotic induction, whereas induced untreated samples displayed a pronounced fibrotic phenotype (Figure 4B).

Treatment with SB-431542, CP-673451 and RepSox effectively suppressed fibrotic marker expression and partially restored a non-fibrotic phenotype. In contrast, AM580 and CCG-1423 showed only partial inhibition of fibrotic remodeling (Figure 4B).

Fibrotic marker expression was quantified by image segmentation and analysis of fibronectin (Figure 4C) and collagen I (Figure 4D) signal intensities. Consistent with qualitative observations, SB-431542 and RepSox showed the strongest inhibition of both markers. AM580 and CP673451 exhibited intermediate effects, with a more pronounced reduction in collagen I than fibronectin, whereas CCG-1423 did not significantly reduce fibronectin or collagen I levels.

In addition to high-content imaging, fibrotic remodeling was assessed by quantifying secreted procollagen I levels in the culture supernatants over time (Figure 4E). Consistent with imaging data, procollagen I was undetectable in control, SB-431542-treated and RepSox-treated conditions. CP-673451and AM580 showed weak and intermediate inhibition, respectively, whereas CCG-1423 did not significantly reduce procollagen I secretion compared with controls (Figure 4F).

## Discussion

Fibrotic remodeling is a central pathogenic mechanism underlying several retinal diseases, including proliferative vitreoretinopathy (PVR), age-related macular degeneration (AMD), and diabetic retinopathy (DR) (Datlibagi et al., 2023). In AMD progression, retinal pigment epithelium dysfunction is widely considered a key driver (Kim et al., 2021) and therefore represents an important target for therapeutic intervention.

Under pathological conditions, RPE cells secrete mediators that recruit and activate inflammatory cells and fibroblasts. Following damage, RPE cells undergo epithelial-to-mesenchymal transition, a process characterized by the loss of apical-basal polarity and the acquisition of a more mesenchymal phenotype. This transition ultimately leads to RPE fibrosis, a hallmark of the disease pathology (Shu et al., 2020).

The in vitro platform presented here consists of human primary fetal RPE cells aggregated to three-dimensional structures and cultured without extrinsic matrix components or artificial substrates. The model is self-organizing and matures as free-floating aggregates. In contrast to Matrigel-dependent organoid cultures and previous studies using primary RPE cells in hydrogels or viscosity-modifying matrices, the hRPE-MT platform generates microtissues under scalable standard culture conditions (Afanasyeva et al., 2021; Eiraku et al., 2011; Sato et al., 2013; Usui et al., 2019).

Microtissues can be generated, cultured, and analyzed in 96-and 384-well plates, making them fully compatible with robotic liquid handling and automated workflows. hRPE-MTs remain stable over several weeks, enabling chronic exposure studies and longitudinal analyses.

The generated spheroids form a distinct peripheral monolayer with a high degree of apical-basal polarity that is stably maintained over extended culture periods. The apical side is oriented toward the culture medium while the basolateral side faces inward. Spheroids exhibit well-defined junctional structures, as demonstrated by characteristic staining of the tight junction marker ZO-1. Furthermore, hRPE-MTs display a strong retinal identity at the transcriptional level, as evidenced by the expression and maturation-dependent induction of hallmark retinal genes. Gene set enrichment analysis over time reveals that the most enriched gene sets are associated with core retinal processes including the visual cycle, RPE specific functions, transmembrane transport, and photoreceptor-related pathways.

Compared to broadly used 2D transwells, the 3D hRPE microtissue model offers a superior biomimetic architecture where cells self-organize into a polarized geometry with native extracellular matrix deposition. The matrix-free 3D environment eliminates artificial membrane contact, promoting sustained epithelial differentiation and cellular longevity over several weeks. Furthermore, aggregating microtissues in standard microplates eliminates the optical interference associated with cell culture inserts, enabling seamless integration with automated liquid handling and high-content imaging. While 2D transwells remain advantageous for specialized endpoints requiring directional fluid collection or transepithelial electrical resistance (TEER) measurements, the 3D spheroid model offers superior scalability and physiological fidelity for high-throughput phenotypic screening.

Previous studies have shown that RPE cells can be induced to undergo EMT through stimulation of key signaling pathways, including TGF-β, IL-6, IL1β and TNFα (Boles et al., 2020; X. Chen et al., 2020; Y. Chen et al., 2021). In line with these findings, hRPE-MTs are highly responsive to co-stimulation with TGF-β2 and TNF-α, which induces robust EMT and fibrotic changes. Upon induction, microtissues undergo pronounced morphological changes, including compaction and loss of the peripheral monolayer structure.

Tight junctions are essential for maintaining cell–cell contacts and regulating solute gradients. Consistent with EMT induction, we observed a loss of ZO-1 localization, indicating disruption of tight junction integrity. Concurrently, tissues begin to express fibrotic markers such as fibronectin, collagen I, and α-smooth muscle actin (α-SMA), reflecting a transition toward a myofibroblast-like phenotype. Increased secretion of pro-collagen I was detected in culture supernatants, and transcriptomic profiling revealed strong upregulation of EMT-associated genes alongside downregulation of epithelial and RPE-specific markers.

To demonstrate the utility of the model, we evaluated five compounds known to inhibit EMT and fibrosis through distinct mechanisms of action. The TGF-β inhibitors SB-431542 and RepSox exhibited the strongest efficacy, markedly reducing the expression of EMT-associated markers. Inhibition of the canonical TGF-β/SMAD pathway is known to block early EMT induction, a response that was recapitulated in the hRPE microtissue model. Treatment partially restored fibrotic marker levels (fibronectin, collagen I, and secreted pro-collagen I) toward baseline and led to partial recovery of the cyst-like morphology observed in untreated tissues.

Previous studies have demonstrated that, in vitro, complete reversal of the fibrotic phenotype in RPE cells typically requires combination treatments (Y. Chen et al., 2021). Accordingly, inhibition of canonical TGF-β/SMAD signaling alone may be insufficient, and multi-target intervention strategies may be necessary to achieve full phenotypic reversion in the hRPT-MT model.

Platelet-derived growth factors (PDGFs) and their receptors (PDGFRs) are critical mediators of pathological fibroblast-like cell survival and are implicated in the stabilization of the fibrotic phenotype following the early stages of EMT. Consistent with this role, treatment with the PDGF antagonist CP6734451 resulted in a partial attenuation of fibrotic marker expression.

Fibronectin remained above 50% of the fibrotic control, whereas collagen I, representing a later-stage marker associated with fibrotic maturation and extracellular matrix deposition, was reduced to a markedly lower level. This differential response is in line with observations from human RPE systems, where PDGF/PDGFR signaling has been shown to predominantly regulate proliferative, migratory, and contractile aspects of RPE activation rather than serving as a primary driver of EMT initiation. In human RPE cells, PDGF isoforms induce PDGFR phosphorylation and promote proliferation and migration, while PDGF-BB specifically enhances chemotactic migration through PDGFRβ-dependent PI3K/Akt and MAPK signaling pathways (Chan et al., 2013; Li et al., 2007). Taken together this supports the interpretation that PDGFR inhibition by CP6734451 primarily interferes with downstream fibrotic maturation and cellular remodeling processes, including extracellular matrix deposition and contractility, rather than fully blocking upstream EMT-inducing signals.

The retinoic acid receptor-α (RAR α) agonist AM580 did not inhibit fibronectin levels but elicited an approximately 40% decrease in collagen I expression. Consistent with prior reports, RAR signaling inhibits EMT in RPE cells primarily through transcriptional regulation rather than direct interference with TGF-β signaling (Kobayashi et al., 2019). Correspondingly, the associated cellular phenotypic changes were observed to occur with delayed kinetics.

CCG-1423 targets the Rho/MRTF/SRF transcriptional axis, a key regulator of cytoskeletal remodeling. Under the conditions tested, CCG-1423 did not significantly alter the expression of collagen I, secreted pro-collagen I, or fibronectin. This suggests that its inhibitory effects likely occur downstream of canonical TGF-β signaling and are more specifically directed toward contractility-and mechanotransduction-related processes rather than broadly suppressing the EMT program.

## Conclusion

Subretinal fibrosis driven by epithelial-mesenchymal transition of retinal pigment epithelial cells remains a major cause of irreversible vision loss in advanced age-related macular degeneration (AMD).

Here we describe the development of a robust, self-organizing three-dimensional primary human RPE microtissue model that effectively bridges the gap between oversimplified 2D cultures and low-throughput, heterogeneous organoid systems. Through synergistic pro-fibrotic and pro-inflammatory stimulation with TGF-β and TNF-α, this platform recapitulates key hallmarks of fibrosis, including the loss of epithelial polarity, marked morphological compaction, and pathological production of extracellular matrix components such as collagen I and fibronectin.

Importantly, the matrix-and hydrogel-free design enables high reproducibility and compatibility with automated workflows. As demonstrated by the successful phenotypic evaluation of targeted pathway modulators, this scalable microtissue system constitutes a powerful and predictive screening platform, with potential to accelerate the discovery and preclinical development of antifibrotic therapies for AMD and related ocular fibrotic diseases.

## Acknowledgements

We thank the ETH Scientific Center for Optical and Electron Microscopy (ScopeM) and in particular Miriam Susanna Lucas and Falk Lucas, for their support with scanning and transmission electron microscopy (SEM and TEM) imaging, as well as for the processing of all electron microscopy data presented in this manuscript.

During the preparation of this work, the authors used Google’s Gemini (Version 3.5 Flash) to improve language clarity and correct grammar errors. After using this tool, the authors reviewed and edited the content as needed and take full responsibility for the final publication.

## Declaration of conflict of interest

TH, AP and DF are employees of InSphero AG, Schlieren, Switzerland, which commercializes the Akura™ platform. CR and PW are employees of F. Hoffmann-La Roche Ltd., Basel,

---

**Supplementary Figure 1.** Concentration-response curves of pathway modulators used for model validation.

**Supplementary Table 1.** Compound information of pathway modulators used for model validation.

