## Supplementary Materials for "A microtissue-based retinal fibrosis platform for drug efficacy testing"

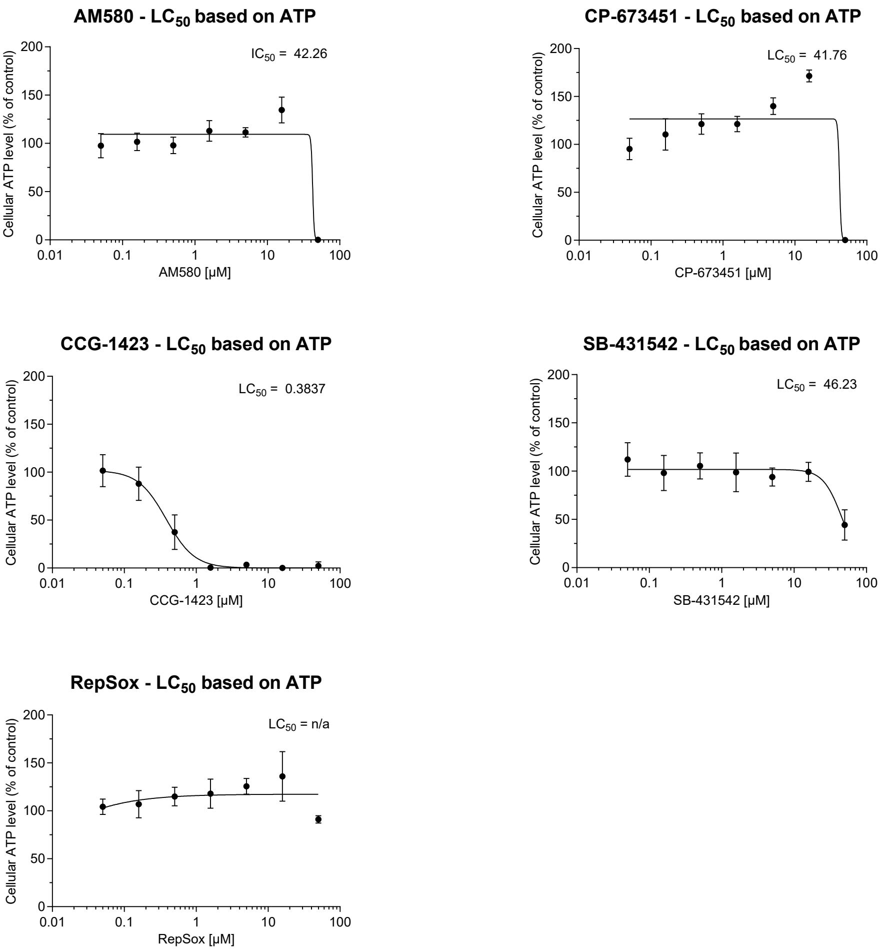


Supplementary Figure 1

| **Compound** | **Supplier** | **Cat. No.** |
|---|---|---|
| SB-431542 | TargetMol | T1726 |
| RepSox | TargetMol | T6337 |
| AM580 | TargetMol | T5854 |
| CCG-1423 | TargetMol | T2014 |
| CP-673451 | TargetMol | T6091 |

Supplementary Table 1
